# Passive endocytosis in model protocells

**DOI:** 10.1101/2023.01.07.522792

**Authors:** Stephanie J. Zhang, Lauren A. Lowe, Palapuravan Anees, Yamuna Krishnan, Thomas G. Fai, Jack W. Szostak, Anna Wang

## Abstract

Semipermeable membranes are a key feature of all living organisms. While specialized membrane transporters in cells can import otherwise impermeable nutrients, the earliest cells would have lacked a mechanism to import nutrients rapidly under nutrient-rich circumstances. Using both experiments and simulations, we find that a process akin to passive endocytosis can be recreated in model primitive cells. Molecules that are too impermeable to be absorbed can be taken up in a matter of seconds in an endocytic vesicle. The internalized cargo can then be slowly released over hours, into the main lumen or putative cytoplasm. This work demonstrates a way by which primitive life could have broken the symmetry of passive permeation prior to the evolution of protein transporters.

## Introduction

In extant life, membranes provide a selective barrier between a cell and its environment, which enables the inheritance of adaptive traits and ultimately leads to Darwinian evolution.^1,2^ Life itself may have emerged from self-replicating informational molecules spatially constrained by primitive membranes.^1–3^ Amongst various lipid candidates, fatty acids are particularly well-suited as the components of primitive membranes owing to their prebiotic relevance^4,5^ and dynamic exchange properties^6,7^. The possible involvement of fatty acid vesicles in the origin of life has been further demonstrated by their ability to grow and divide without complex biochemical machinery^8–10^ and to encapsulate RNA templates that are being non-enzymatically copied.^11,12^

In nascent life, the absence of protein transporters implies that a protocell enveloped by lipid bilayers would have had to rely on passive diffusion for internalizing nutrients.^13–15^ Diffusive transport across the membrane is symmetric and is driven purely by the concentration gradient of solutes across the membrane. Despite this, the fluxes need not be the same. When the symmetry is broken, for example, by placing a nutrient sink within the cell, or by a changing external environment, there can be a net flux into or out of the protocell. However, under transient nutrient-rich circumstances, the low permeability of primitive membranes, needed to prevent the loss of the encapsulated cargo, also delays the efficient acquisition of nutrients. Hence, a protocell would have to reside within a pool of nutrients for hours to absorb useful levels of polar or charged molecules.^12,15^ Whether physicochemical stimuli can fuel non-diffusive transport mechanisms remains an important question because such mechanisms could break the trans-membrane symmetry in a way that could bias the inward nutrient flow, or expedite nutrient import.

One type of transport that has both ‘active’ and ‘passive’ modes is endocytosis. In modern biology, the inward budding of lipid membranes can be active such as in receptor-mediated endocytosis, or passive as in fluid-phase endocytosis. Such a higher-order topological transformation has been previously demonstrated in phospholipid membranes via various pathways.^16–18^ However, the ability of model primitive membranes to endocytose, i.e., internalize cargo from the external milieu, has yet to be definitively shown. Model primitive membranes typically consist of lipids with dynamic properties distinct from those of phospholipids and thus the routes to engender shape changes could potentially differ^19^. For instance, flip-flop of model primitive membrane lipids rapidly relaxes any curvature stress^20^ that might otherwise help drive the shape transformation and help overcome the energy barrier required for a topological change^19^. On the other hand, flip-flop may also be useful for enabling the membrane to adopt the extreme configurations required for inward or outward budding.

Here, we demonstrate that primitive cell compartments composed of fatty acids can passively endocytose via a purely physicochemical process. A simultaneous reduction in volume and increase in surface area allows larger molecules to be imported via an inward bud into model protocells, mimicking the process of passive endocytosis in complex eukaryotes. This process takes only seconds, similar to modern forms of endocytosis. The protocells continue to retain the internalized solutes even when removed from the nutrient source, and are thereby provided time to absorb the nutrients. Further, we construct a numerical model that captures such out-of-equilibrium shape changes. Taken together, we believe this marks an important step toward understanding the development of solute internalization and transport in primitive cells.

## Results and Discussions

### Fatty acid vesicles undergo membrane invagination and internal budding

Here we study the shape transformations and subsequent topological transitions of fatty acid vesicles in response to volume and area perturbations. For a sphere to change shape, it must have an increase in surface area to volume ratio. In principle, this can be accomplished via two pathways: by reducing the volume or increasing the surface area. We explore shape changes obtained through combinations of these pathways. We use hyperosmotic shocks to draw water out of the vesicles and reduce the internal volume; the magnitude of the osmotic shock is denoted as a change in concentration of the salt (Na-bicine), ΔC_V_, from its starting point of 50 mM. We achieve membrane area growth by adding alkaline oleate micelles to a buffered solution of vesicles; the increase in total membrane lipid is denoted as a change in lipid concentration, ΔC_A_, relative to a starting lipid concentration of 0.5 mM. Evaluations of the vesicle radius before and after micelle addition suggest that the increase in surface area is no more than two-fold under the conditions used in this work (see SI). To observe shape changes in real-time with fluorescence microscopy, we prepared oleic acid/oleate (C18:1) giant unilamellar vesicles (GUVs) encapsulating water-soluble and membrane-impermeable fluorescent dyes (either 0.5 mM HPTS 8-hydroxypyrene-1,3,6-trisulfonic acid trisodium salt, or 1 mM calcein blue). These vesicles were then diluted 1:10 in buffer that was identical in composition and pH to the original buffer to provide good contrast between encapsulated and unencapsulated dyes, enabling visualization of the dye encapsulated within vesicles.

We previously reported that an increase in lipid concentration by micelle addition (ΔC_A_ = 1 mM) to oleate GUVs results in outward budding, with the GUVs transforming into a series of smaller vesicles in close proximity, and possibly still connected by tethers (Figure 1(a)).^9^ Addition of a higher concentration of micelles (ΔC_A_ > 1 mM) resulted in vesicle division. In terms of a putative geological setting for such an event, fluid flow from precipitation, in pools, or from tectonic or volcanic events that drive hydrothermal processes, might have carried and mixed nutrients, mineral particulates, and lipid components. Such episodic events from fluid flow-induced geochemical processes might result in fluctuations in osmotic pressures and simultaneous release of nutrients and lipids. Because increasing the surface area and reducing the internal volume would be predicted to increase the surface area to volume ratio, we expected that doing both simultaneously, denoted as stimulus A (ΔC_V_ = 100 mM, ΔC_A_ = 1 mM), might also cause vesicle division.

**Figure 1.**
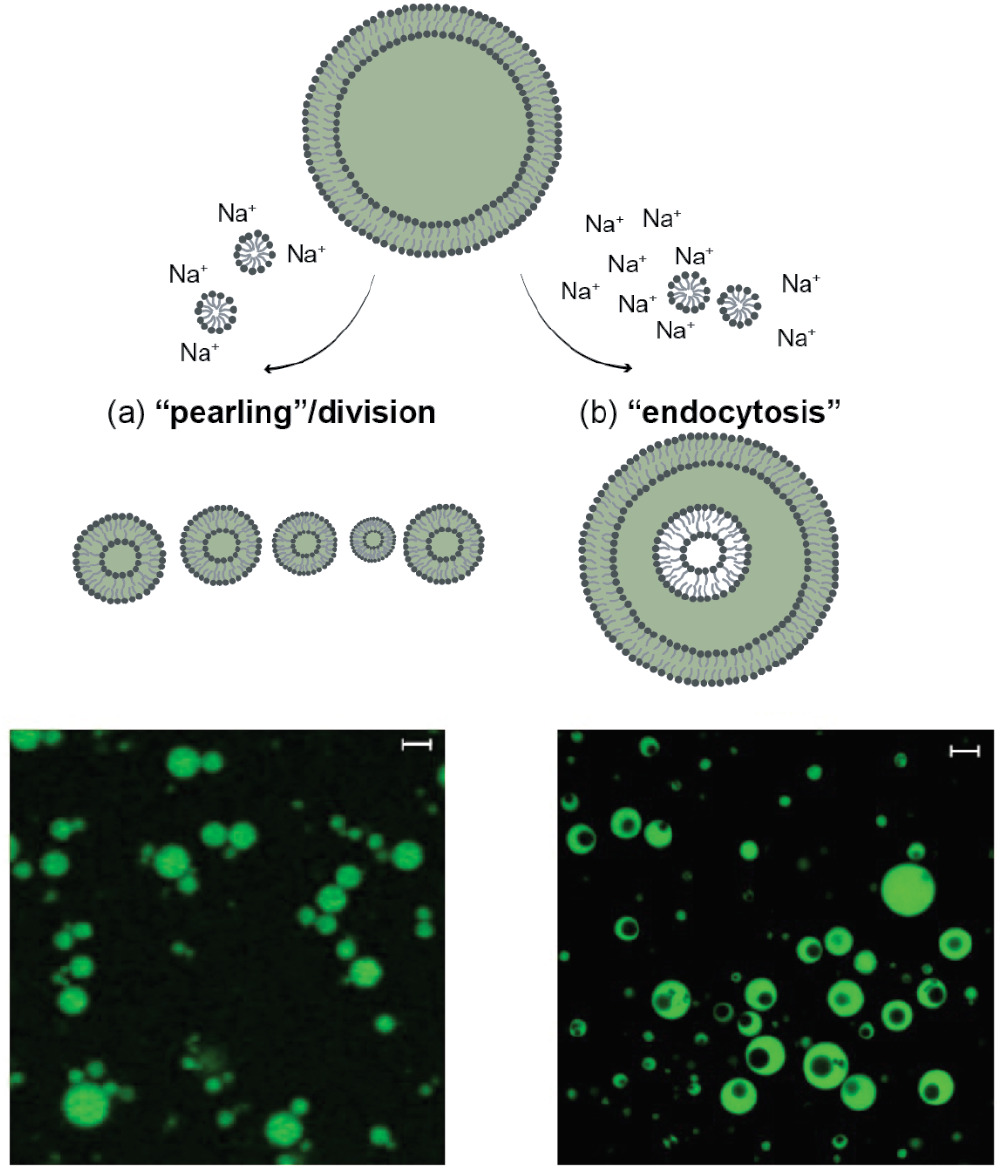
Schematics and confocal micrographs of topological transformations of oleic acid/oleate giant unilamellar vesicles (GUVs) following addition of oleate micelles or exposure to hyperosmotic shock. a) Outward budding of GUVs was induced by micelle addition (stimulus B; Δ*C*_A_ = 1 mM). The resulting smaller, closely-spaced vesicles can be seen in the confocal micrographs. This is similar to the supplemental videos shown in our previous work.^9^ b) GUVs could be induced to “endocytose” upon a hyperosmotic shock coupled with micelle addition (stimulus A; ΔC_V_ = 100 mM, ΔC_A_ = 1 mM. See also Movie S1). Confocal micrographs reveal vesicles initially containing 0.05 M Na-bicine and labeled with 0.5 mM HPTS (green fluorescence) have compartments that appeared as dark voids. Scale bars represent 5 μm.

Instead, a series of different dramatic shape transformations occurred upon a simultaneous introduction of membrane material and an osmotic shock (Figure 1(b), Figure S1, and Movies S1-S8; please see Methods for exact experimental conditions). Specifically, GUVs appeared to elongate, then rapidly transform first into a stomatocyte via invagination and then into a sphere. We confirmed that the topological transformation was to a vesicle-in-vesicle structure rather than a torus by taking sectional image slices along the z-axis and reconstructing the volume in 3D (Movie S2). Further, we performed two distinct experiments to show that the internal vesicles are indeed fully separated from the external membrane. First, we added an aqueous dye, Alexa 594 hydrazide, that is exclusively aqueous and does not bind to membranes, to a final concentration of 10 μM. Alexa 594 hydrazide stayed completely outside of the GUV and did not enter the interior of the endocytosed vesicles, which demonstrates that the inner vesicles are not connected to the outside (Figure S2). Second, we added the dye Voltair^PM 21^(final concentration: 250 nM) which localizes to the fatty acid membrane by its POPE (1-palmitoyl-2-oleoyl-sn-glycero-3-phosphoethanolamine) moiety, but cannot flip-flop across the external membrane or even reach any inner compartment owing to its conjugation to polar DNA oligonucleotides. The observation that only the GUV outer membrane leaflet became labeled excludes the possibility that the inner vesicle membranes are connected to the outer GUV membrane (Figure S3). Therefore, we can conclude that instead of budding *outward* (stimulus B; ΔC_V_ = 0 mM, ΔC_A_ = 1 mM Figure 1(b)), the vesicles had budded *inward*. We hypothesize that the complete budding of an internal vesicle was possible because both the generated lateral tension from the insertion of lipids into the outer leaflet, and the curvature stress from flip-flop not transporting lipids into the inner leaflet as rapidly as lipids inserting into the outer leaflet, were sufficient to overcome the energies required for breaking the neck and completing the budding process.^22^ The exact mechanism is still uncertain, and should be the subject of future work.

### Protocells take up cargo passively following membrane invagination

As previously mentioned, micelle addition by itself leads only to outward budding^8,9^. Interestingly, micelle addition (ΔC_A_ = 0.5 mM to 5 mM) accompanied by varying levels of osmotic shock (ΔC_V_ = 25 mM to 100 mM), consistently led to inward budding. By exploring this two-dimensional parameter space, we found that the efficiency of internal compartment formation increased with the magnitude of the surface area increase. As more micelles were added (ΔC_A_ = 0.5 mM to 5 mM), the fraction of vesicles with internal compartments increased, and this effect was evident across all magnitudes of osmotic shock (ΔC_V_ = 25 to 100 mM, Figure 2). For example, the population of vesicles with internal compartments increased from approximately 7.3 ± 1.3 % (ΔC_A_ = 0.5 mM; ΔC_V_ = 100 mM, total vesicles analyzed = 3840) to approximately 88.5 ± 3.0 % (ΔC_A_ = 5 mM; ΔC_V_ = 100 mM, total vesicles analyzed = 1169).

**Figure 2.**
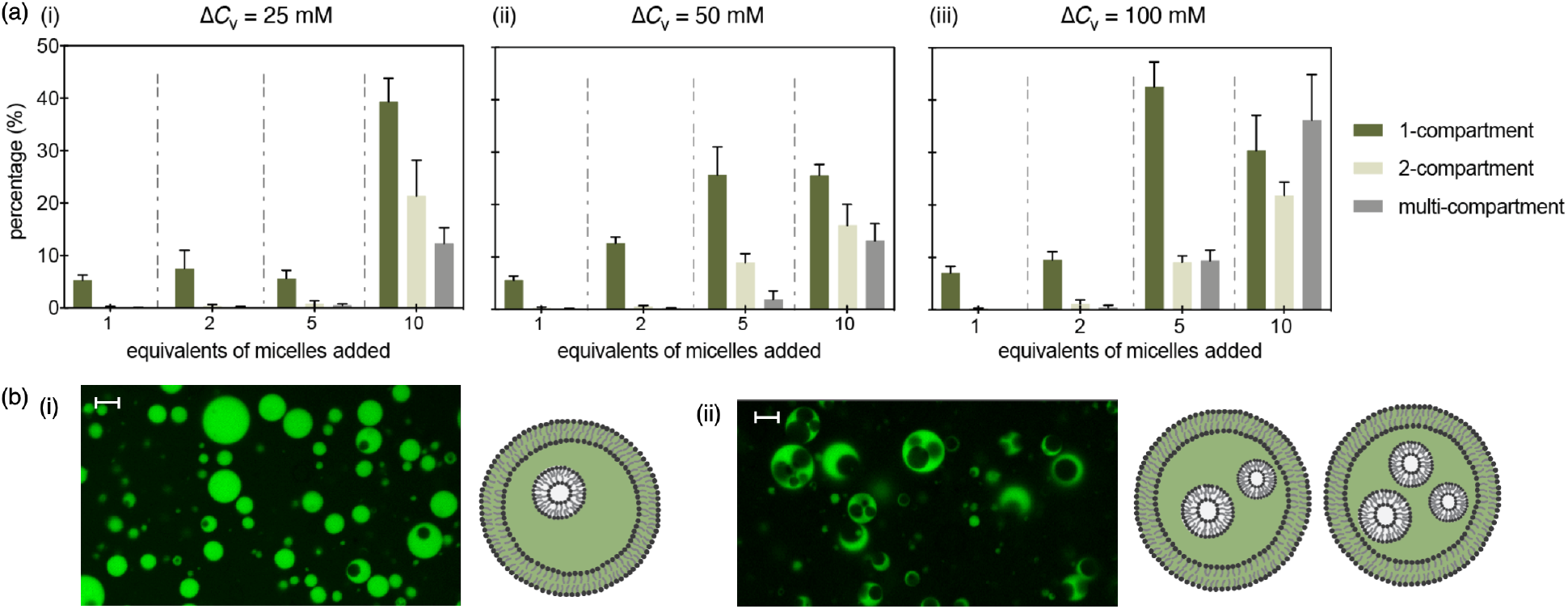
Number of compartments per vesicle as a function of osmotic shock and micelle addition. Conditions are summarized in Table 1. (a) Distributions of compartments per vesicle for osmotic shocks and micelle additions of different magnitudes (Δ*C*_V_ = 25 (i), 50 (ii), 100 (iii) mM Na-bicine, Δ*C*_A_ = 0.5, 1, 2.5, 5 mM oleate). The equivalents of oleate in the micellar solution added are defined with respect to the concentration of oleate/oleic acid in the starting vesicle suspension. (b) Representative confocal micrographs of vesicles with (i) single and (ii) multiple internal compartments. Scale bars represent 5 μm. Each condition consisted of at least 5 replicates, error bars indicate the standard deviation from the mean. The total number of analyzed vesicles for each condition is indicated in Table 1.

We also found that the number of internal compartments could be tuned by varying the magnitude of the osmotic shock. At concentrations of added micelles between 2.5 mM to 5 mM, increasing the osmotic shock increased both the number of vesicles with compartments, and the occurrence of multi-compartment vesicles. For example, at an added micelle concentration of 5 mM, the fraction of vesicles with internal compartments increased from approximately 70.1 ± 9.9 % (at ΔC_V_ = 25 mM, total vesicles analyzed = 695) to approximately 88.5 ± 3.0 % (at ΔC_V_ = 100 mM, total vesicles analyzed = 1169), while the fraction of multi-compartment vesicles increased 3-fold from approximately 12.3 ± 3.0 % (ΔC_V_ = 25 mM, total vesicles analyzed = 695) to approximately 36.2 ± 8.7 % (ΔC_V_ = 100 mM, total vesicles analyzed = 1169).

Creating two- or multi-compartment vesicles required a higher concentration of added micelles for lower osmotic shocks: A non-negligible yield of two-compartment or multi-compartment vesicles required ΔC_A_ = 5 mM at a lower osmotic shock (ΔC_V_ = 25 mM), whereas it required only ΔC_A_ = 2.5 mM at a higher osmotic shock (ΔC_V_ = 100 mM). At lower concentrations of added micelles (ΔC_A_ = 0.5 mM to 1 mM), varying the magnitude of osmotic shock (ΔC_V_ = 25 mM to 100 mM) did not lead to significant changes in the number of internal compartments formed.

Having determined that the inward budding was robust, we then tested whether this pathway allowed for the passage of large polar molecules into protocells prior to the evolution of complex protein machinery. We found that passive endocytosis allowed the long DNA-dye conjugate (5′-Cy5-C_(10)_ A_(18)_-3′) (Figure 3 (a)) to be imported from the external medium into the vesicle itself. Specifically, the Cy5-labeled DNA 28-mers (appearing magenta) were introduced along with the simultaneous osmotic shock and micelle addition (stimulus A) into oleic acid/oleate GUVs containing encapsulated calcein blue (appearing blue). The resultant formation of magenta interior compartments indicates that the Cy5-labeled DNA 28-mers were successfully transported to the lumen of the internalized vesicles and thereby internalized by the GUVs (Figure 3(b)(i,ii)).

**Figure 3.**
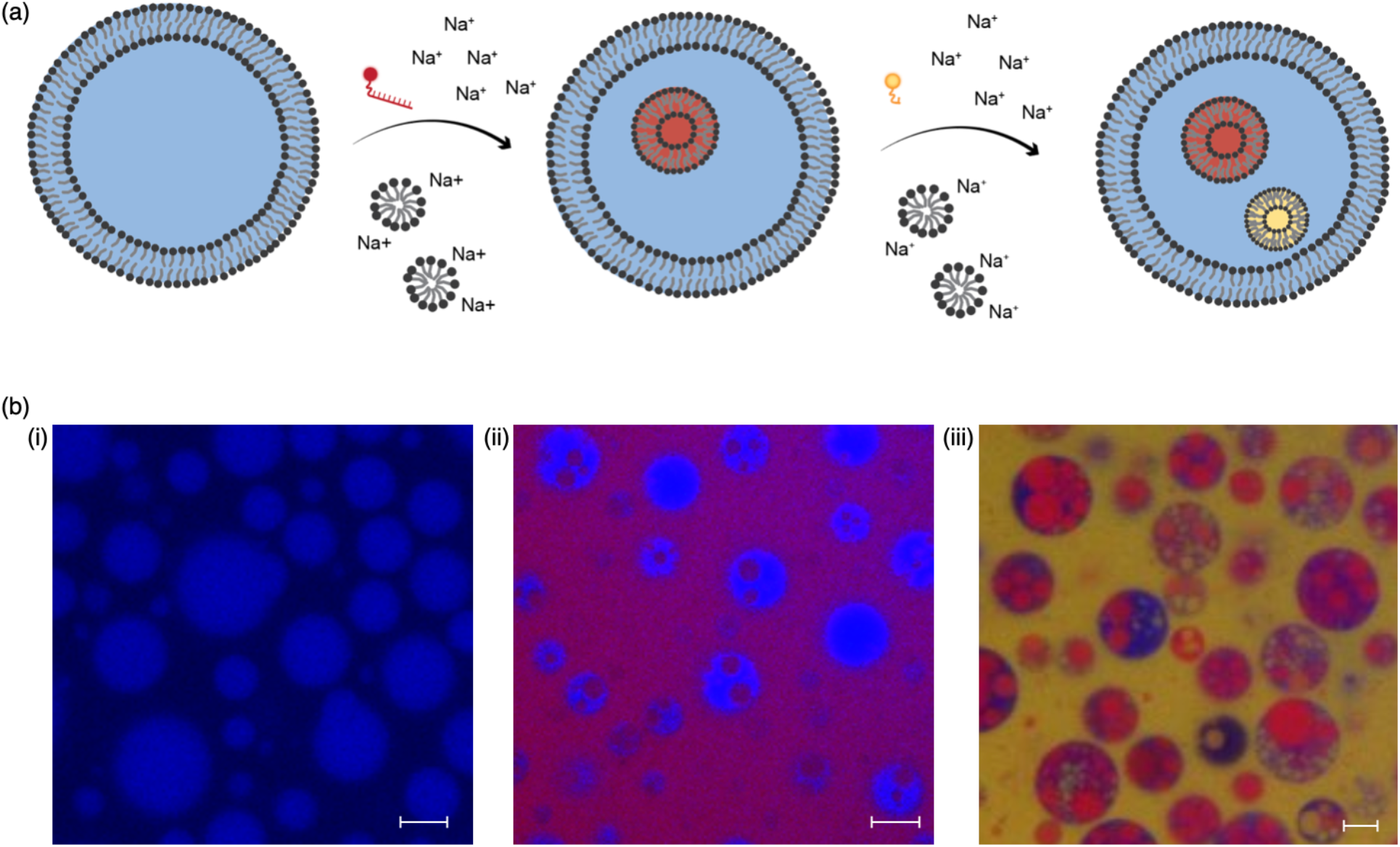
Induction of multiple sequential passive endocytosis events (a) Schematic for induction of multiple passive endocytosis events induced by successive osmotic shocks coupled with micelle additions. (b) Confocal micrographs of fatty acid vesicles with various numbers and sizes of interior compartments. (i) Initial GUVs with encapsulated calcein blue (blue). (ii) Simultaneous addition of 1 μM 5’-Cy5-C_(10)_ A_(18)_-3’, ΔC_V_ = 100 mM Na-bicine, and ΔC_A_ = 2.5 mM oleate, resulted in inner compartments containing encapsulated 5’-Cy5-C_(10)_ A_(18)_-3’ (magenta) derived from the external solution. (iii) A second round of passive endocytosis stimulated by the simultaneous addition of 5 μM Fluorescein-12-UTP, ΔC_V_ = 100 mM Na-bicine, and ΔC_A_ = 2.5 mM oleate, resulted in new inner compartments containing encapsulated Fluorescein-12-UTP (yellow) derived from the external solution. Scale bars represent 5 μm. The introduction of multiple heterogeneous stimuli to a uniform starting solution of vesicles ultimately leads to the emergence of population diversity (see also Movie S12).

The resultant vesicle-in-vesicle structures were not expected to preclude further inward budding events. We therefore tested whether repeated inward budding could be induced by successive rounds of simultaneous osmotic shock and micelle addition (stimulus A). Following the intake of Cy5-labeled DNA 28-mers into one set of internalized buds, a second simultaneous osmotic shock and micelle addition event was induced in the presence of the yellow aqueous compound fluorescein-12-UTP in the exterior solution. This resulted in the UTP being internalized via a second set of buds: the blue vesicles contained both magenta and yellow inner compartments (Figure 3(b)(iii)). With both stimulus events being in the regime where two or more internal compartments are likely (Figure 2), the resultant vesicles displayed a high internal volume fraction of internalized vesicles. The excess micelles introduced also led to *de novo* vesicle formation: magenta vesicles containing yellow inner compartments are also occasionally observed. Some compartments also appear intermediate in color (e.g. orange), suggesting some mixing between compartments might have occurred.

This result is informative for two reasons. First, the presence of inner compartments that are a different color from the external medium (after two cycles of micelle addition in the presence of different dyes) indicates that the shape transformation of the vesicles does not end in the stage of stomatocytes (Figure S1(a)). Rather, inward vesiculation was completed and a full topological transformation had occurred (Figure S1(b)). Without the final topological transformation, all the endocytic compartments would be the same color as the external medium. Second, the presence of both yellow and magenta internal compartments is consistent with two separate rounds of endocytosis having taken place. This confirms that the simultaneous osmotic shock and micelle addition event can trigger repeated endocytic events.

### Release of cargo from internal compartment into main lumen

A substantial uptake of nutrients from the external solution by passive diffusion across the limiting membrane of protocells requires them to reside for hours within a pool of nutrients. By contrast, passive endocytosis results in a net inward flow of cargo within seconds, although the cargo is still contained within a membrane. Releasing nutrients stored inside the endocytic compartments into the main lumen i.e. putative cytoplasm would allow them to interact with other components within the protocell and is the final critical step in this pathway for nutrient uptake. For example, this could enable dinucleotides to interact with replicating internal oligonucleotides.

To test the ability of the internal endocytic compartment to release nutrients, we first used a relatively impermeable substrate, the fluorescein-labeled cyclic dinucleotide (cGAMPfluo) as a fluorescent tracer. cGAMPfluo was introduced along with a simultaneous osmotic shock and micelle addition (ΔC_V_ = 100 mM, ΔC_A_ = 1 mM) to oleic acid/oleate GUVs (Figure 4(a)). The vesicles were then diluted five-fold into a 200 mM glucose buffer that was identical in composition and pH to the original buffer but lacked the lipids and encapsulated solute, to dilute the free cGAMPfluo in the external medium. We then took confocal microscopy images over time to measure the fluorescence intensities in the internalized compartments, the main lumen of the GUVs, and the external medium. The fluorescence intensity serves as a measure of the relative concentrations of encapsulated material.

**Figure 4.**
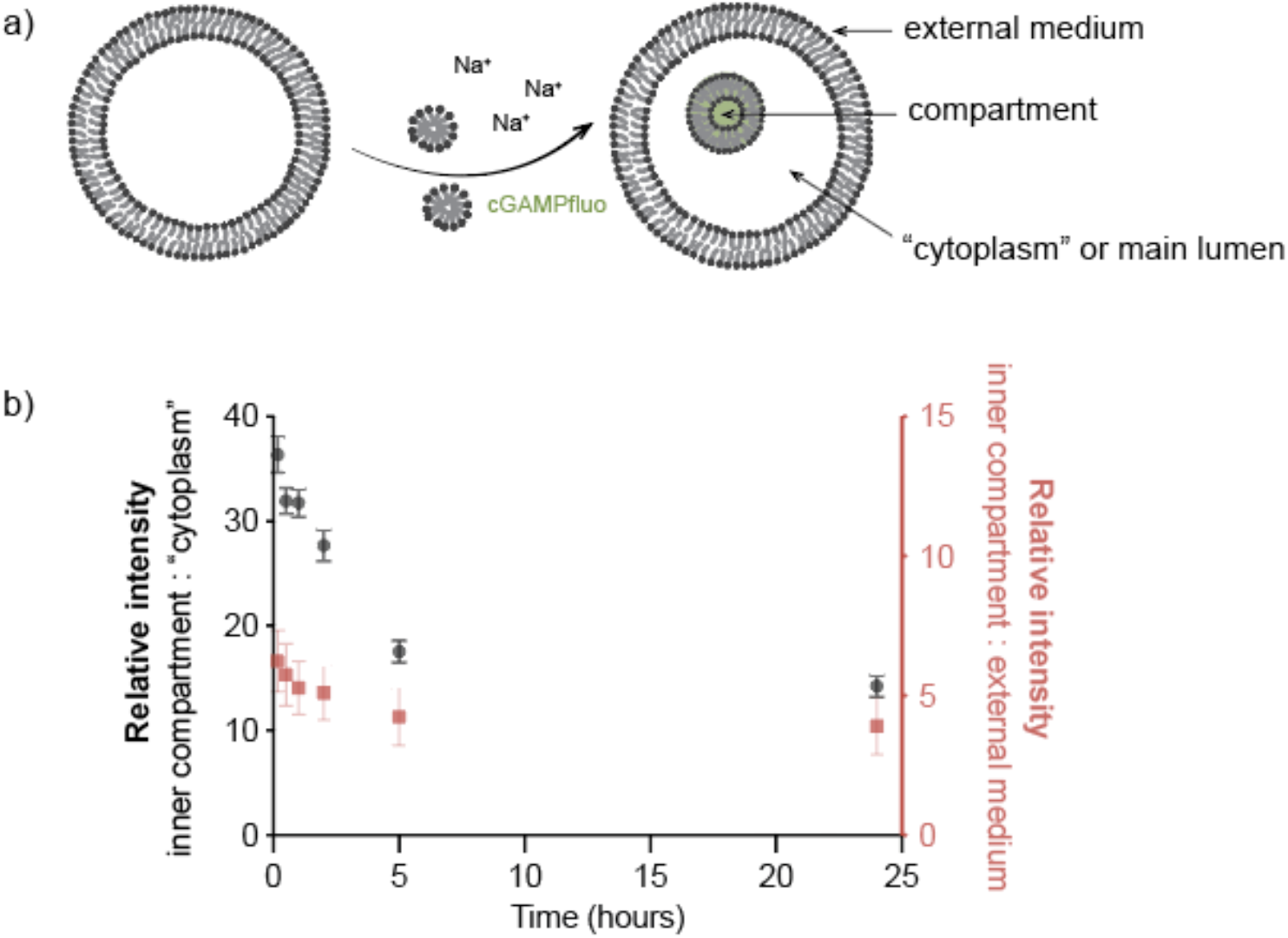
Internal compartments can slowly deliver nutrients of low permeability to the main lumen or ‘cytoplasm’ of a protocell. (a) Schematic of fluorescein-labeled cyclic dinucleotide (cGAMPfluo), a relatively impermeable substrate, being encapsulated in the interior compartment via one hyperosmotic shock coupled with micelle addition (Δ*C*_V_ = 100 mM Na-bicine, Δ*C*_A_ = 2.5 mM oleate, *C*_cGAMPfluo_ = 0.1 mM). (b) The ratio of fluorescence intensity of the internal compartment vs the ‘cytoplasm’ decreases with time; similarly, the ratio of fluorescence intensity of the internal compartment vs the external medium decreases with time, indicating that the dinucleotide inside the internal compartment is slowly entering the ‘cytoplasm’.

Tracking the fluorescence intensity of the internalized compartments relative to the external medium (Figure 4(a)), which we consider an infinite bath, showed a decrease from 6.2 ± 1.0 to 3.9 ± .9 (N=124) over 24 hours. This suggests that the oligomer was slowly released by the internalized compartment, into the main lumen. Indeed, the fluorescence intensity of the internalized compartments relative to the main lumen decreased more than two-fold, from the ratio of 36.3 ± 1.7 to 14.3 ± 1.1 over 24 hours (Figure 4(b), N=113). This indicates that the oligonucleotide was slowly released from the internalized compartment into the main lumen. We found similar results for another representative molecule with low permeability to fatty acid membranes,^23^ HPTS (Figure S5). Overall, these results demonstrate that internalized compartments can be used as nutrient storage depots, which would slowly release contents to the main lumen or ‘cytoplasm’ of a protocell. Overall, our studies show that protocells are capable of a primitive form of endocytosis consisting of three characteristic steps: membrane invagination and budding, uptake of soluble external cargo into the resultant internalized compartments and cargo release from the compartments into the main lumen. These constitute the three basic steps of all forms of modern endocytosis.

### Modeling out-of-equilibrium topological changes

To understand the reason for invagination, we model the vesicle as an inextensible elastic material with prescribed bending modulus, spontaneous curvature, and permeability. While the model can not predict whether a full topological transformation can occur, it does enable the prediction of morphological changes preceding a potential topological transition. The surrounding fluid is considered as an aqueous solution that contains amphiphiles that incorporate into the vesicle membrane at a given rate. As shown in Ruiz-Herrero *et al*.,^19^ the resulting behavior may be described in terms of two dimensionless parameters that govern the steady-state vesicle morphology. In addition, to capture the effect of osmotic shocks, we include the possibility of osmotic pressures that drive flows across the membrane. This is done by adding an appropriate normal force to the membrane surface corresponding to the van’t Hoff pressure *p* = *ck*_B_*T*, where *c* is the difference in osmotic concentrations across the membrane.

After applying osmotic shocks in the presence of permeation and membrane growth, we observe that vesicles develop stomatocyte morphologies that progress spontaneously to develop inward buds (Figure 5(a,b)). The initial hyper-osmotic shock causes the vesicles to shrink in volume while adding additional surface area. This increases the surface area to volume ratio over time. We use parameters listed in Table 2 and confirm that the timescales of volume loss and area increase are consistent with those of prior work.^24^

**Figure 5.**
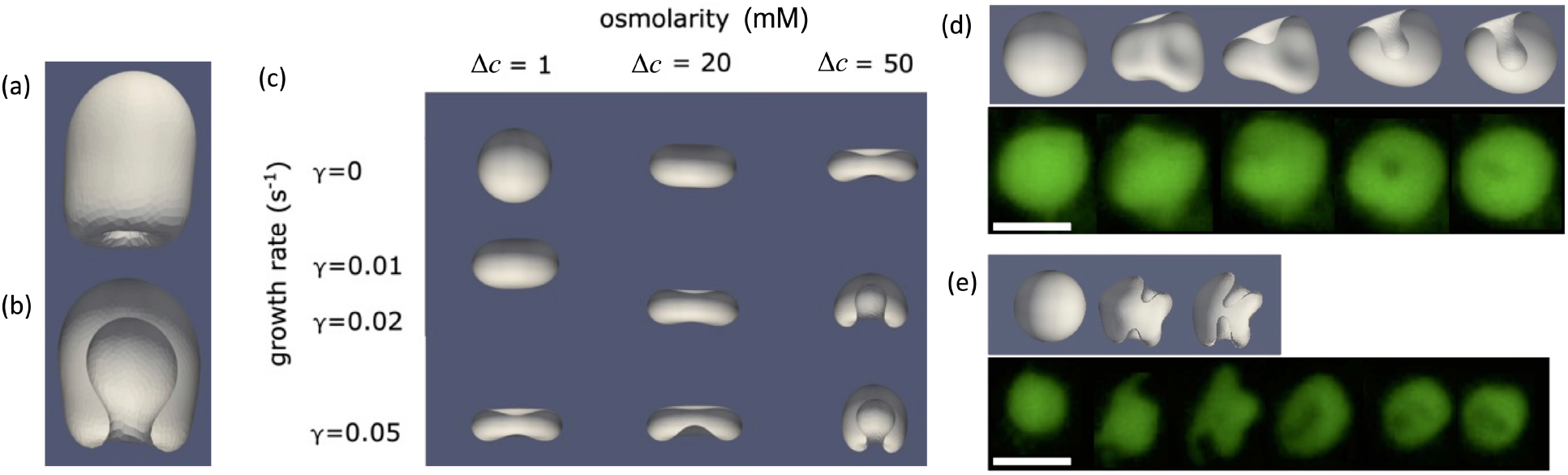
Modeling of vesicle topological changes. For parameters used see Table 2. (a) Side view and (b) cutaway of the final morphology of an initially spherical vesicle after 2 s of growth, with spontaneous formation of an incomplete inner bud, yet to be pinched off. (c) Final morphology of an initially spherical vesicle after 3 s of growth, for different values of the growth rate (γ) and osmolarity (Δ*c = c*_out_ - *c*_in_). Increasing the osmotic shock results in a higher effective growth rate, thereby reducing the threshold for vesiculation. (d) Comparison between simulations (gray) and experiments (green) showing inward vesiculation for a single compartment and (e) two compartments. Scale bars represent 5 μm. Further examples of inward vesiculation in simulations and experiments can be found in Movies S3-S11.

**Table 1:** Summary of different stimuli used to trigger shape changes in vesicles.

| Volume(Na-bicine); Volume(micelles) (μL);<br>*total number of vesicles analyzed* | Micelles (relative to the fatty acid concentration) | | | |
| --- | --- | --- | --- | --- |
| Osmotic shock [Na-bicine] (mM) | 1<br>equivalent<br>(eqv.) | 2 eqv. | 5 eqv. | 10 eqv. |
| 25 | 2.7; 0.5;<br>4800 | 2.7; 1;<br>4269 | 2.7; 2.5;<br>3786 | 2.7; 5;<br>695 |
| 50 | 5.56; 0.5;<br>4395 | 5.56; 1;<br>5897 | 5.56; 2.5;<br>1730 | 5.56; 5;<br>1576 |
| 100 | 11.76; 0.5;<br>3840 | 11.76; 1;<br>2941 | 11.76; 2.5;<br>1697 | 11.76; 5;<br>1169 |

**Table 2:**
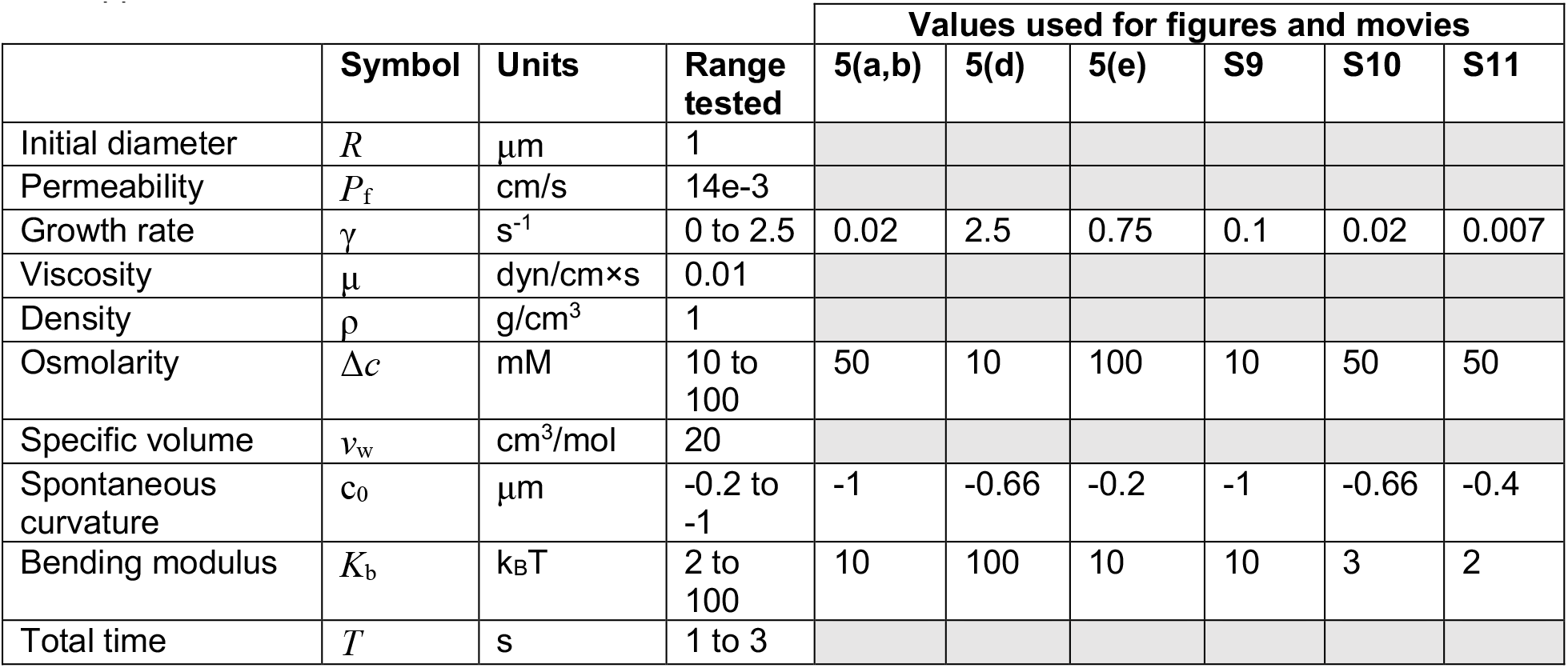
Parameters used in simulations of growing permeable vesicles under osmotic shock.

In terms of the morphological phase plane described in Ruiz-Herrero *et al*.,^19^ the experimental parameter regime corresponds to invagination. We find that the added effect of osmotic pressures is parallel to a higher effective membrane growth rate and lowers the threshold for vesiculation (Figure 5(c)). Our model recapitulates several of the behaviors observed experimentally, including the spontaneous onset of inward vesiculation on the correct timescale of seconds. In terms of the resulting vesicle morphology, we show that the resulting effect from a hyperosmotic shock corresponds to that of an increased effective membrane growth rate. This rationale is consistent with the progressively lower vesiculation threshold that is observed in experiments on vesicles subjected to increasing osmotic shocks (Figure 2).

Furthermore, we find that reducing the magnitude of the spontaneous curvature and the bending modulus can lead to several internal invaginations forming (Figure 5(e), Movies S10-S11). These results suggest that the addition of micelles and an osmotic shock may both contribute towards decreasing the bending modulus of the membrane. Indeed, this is consistent with our recent results showing that either a slight increase in pH or the addition of salt can decrease the bending modulus of fatty acid membranes.^9^

In general, the time series of modeled geometries (Movies S9-S11) correspond well to experimental video microscopy data showing the endocytosis process (Movies S3-S8, see Methods). The changes in shape shown in the movies and in the sequences of images illustrated in Figures 5(d) and 5(e) take place on the timescale of seconds for both experiments and simulations. The relative timescales of water efflux, which is expected to be rapid, and micelle-driven surface area growth (see SI), which is slower, indicate that water efflux and a modest increase in total area are adequate to drive endocytosis within seconds.

Several open questions remain. First, positive spontaneous curvature favors outward budding, whereas negative spontaneous curvature favors inward budding. This suggests that negative spontaneous curvature is important for passive endocytosis to occur, yet the mechanism of how negative spontaneous curvature would arise in our system remains unclear. One possibility is that the bilayer leaflets have an asymmetric interaction with water molecules or ions during simultaneous water efflux and membrane growth. Another hypothesis is that increased external salt bridges the carboxylate headgroups, either decreasing effective membrane area or perhaps changing effective lipid shape to favor negative curvature. Ultimately, while coarse-grained models are effective for predicting shape changes, molecular dynamics simulations of membrane packing might be necessary to reveal the underlying mechanism and cause.^25^ Second, although here we do explicitly solve for the hydrodynamics of the incompressible surrounding fluid, we have not studied the role of hydrodynamics in controlling the vesiculation behavior in detail. More specifically, systematically varying the membrane’s spontaneous curvature and the bending modulus would be an interesting direction for future study.

### Comparison to passive permeability

In light of our experimental demonstration of the release of both dinucleotides and HPTS from the endocytic compartment into the lumen, we also seek to further understand the utility of endocytosis as an inward-transport mechanism and estimate the time that is required for the vesicle’s lumen to obtain various nutrients by passive diffusion across the compartment membrane. Such time scales can be estimated from the net flux (denoted by *J*, with units of number of molecules crossing unit area per unit time). The spontaneous transport of molecules along their concentration gradient across membranes is described by a simple equation

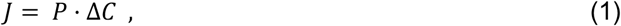

where *P* is the permeability coefficient and Δ*C* (*C*_in_ − *C*_out_) is the concentration difference across the membrane. The number of molecules that cross a given area per unit time can be determined by rearranging Eqn. 1 and the definition of flux

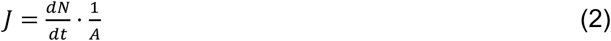

Into

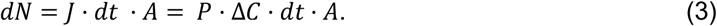

Membrane permeability coefficients (*P*) for short oligomers up to the size of trinucleotides crossing oleic acid membranes, have been previously measured.^12^ We consider trinucleotides, which can enhance non-enzymatic primer extension. The permeability of the oleic acid membrane to an average trinucleotide is *P* ≈ 0.21 . 10 ^−12^ cm/s. In the presence of a 1 mM trinucleotide ‘pool’, the number of trinucleotides that enter a GUV with diameter of 4 *μ*m from the external medium can be estimated from Equation (3) giving 0.06 trinucleotides per second. In other words, it takes approximately 16 seconds for a single trinucleotide to cross the oleic acid membranes when vesicles are surrounded by 1 mM trinucleotides.

We assessed the flux for a range of permeabilities (Figure S6) spanning values for common nutrients (10^−10^ and 10^−13^ cm/s). We found that it takes anywhere between 0.03-33 seconds for a single molecule to cross 4-*μ*m-diameter oleic acid vesicles. It is clear that a protocell would have to reside within such a pool of nutrients for minutes to hours to acquire significant quantities of nutrients by passive diffusion.

This result is in stark contrast to passive endocytosis, which transports a parcel of nutrients inwards within seconds. In this scenario, a dramatic release event is not necessary for the nutrients to reach the putative cytoplasm. Instead, the molecules in the inner compartment can be slowly released into the main lumen of the protocell, where they can then interact with other components. For an inner compartment ¼ the diameter of a 4-*μ*m-diameter GUV, 90% of molecules with permeability comparable to trinucleotides can be released into the lumen within 11 hours (Figure S6(a)). This indicates that protocells that endocytose can absorb a substantial amount of nutrients from a pool despite interacting with it only briefly. Examples include a surface-immobilized protocell capturing nutrients from intermittently nutrient-rich streams of water, or a protocell capturing the released contents from a nearby burst protocell that would otherwise diffuse away.

This principle can be extended further to consider the transport of even larger oligomers. While permeabilities of longer oligomers through oleic acid membranes remain unknown, they are expected to have a lower permeability than trinucleotides. After approximately 4.7 hours (Figure S6(b)), 10% of the molecules with a permeability of 10^−13^ cm/s are released from the interior compartment, with full release within 5 days. This result points to a remarkable function of passive endocytosis – a means of importing otherwise ‘membrane-impermeant’ molecules.

## Conclusions

We have shown that model primitive cell membranes are capable of invagination and inward vesiculation, leading to a complete topological transition to a vesicle-in-vesicle morphology. The number of internal compartments can be controlled by the rate of surface area growth, with increasingly strong osmotic shocks decreasing the rate of surface area growth necessary for vesiculation. We also recapitulated the main results and the relevant timescales in an out-of-equilibrium numerical model. We then found that such inward vesiculation events could lead to internalization of nutrient solutes including mononucleotides and oligonucleotides, drawing further parallels to endocytosis. Such processes could have helped primitive cells capture nutrients that are otherwise impermeable, and could have also generated population diversity from a uniform starting solution of vesicles (Movie S12).

## Acknowledgements

YK acknowledges the support from DP1GM149751, 1R01NS112139-01A1, and Ono Pharma Foundation. TGF acknowledges the support from NSF MCB-2213583 and DMS-1913093. JWS is an Investigator of the Howard Hughes Medical Institute. This work was funded in part by a grant from the Simons Foundation (290363) to JWS. AW acknowledges support from the Australian Research Council (DE210100291), and the Human Frontier Science Program (RPG0029/2020 to AW).

## Materials and Methods

### Materials

Oleic acid (C18:1) was purchased from NuChek Prep (Elysian, MN). DNA labeled with a fluorescent dye (5′-Cy5-C_(10)_ A_(18)_-3′) was synthesized by IDT (Coralville, IA). All other chemicals were purchased from Sigma-Aldrich (St. Louis, MO) and were used without any further purification. Data analysis was performed using ImageJ (version 1.53a), Python, and GraphPad Prism (version 8.4.0).

### Preparation of Micelles and Giant Unilamellar Vesicles (GUVs)

100 mM oleate micelles were prepared by dissolving 50 μmol of neat oleic acid oil in 1 equivalent of NaOH solution to a volume of 500 μL with Millipore water (18.2 MΩ•cm). Fatty acid GUVs were made by resuspending the micelle solution in buffer stock (1 M Na-bicine, pH 8.45), sucrose stock, and Millipore water to the final concentration of 5 mM oleic acid, 50 mM Na-bicine buffer, and 200 mM sucrose as described previously.^9^ Encapsulation of fluorescent dyes was achieved by mixing the solute with the resuspension buffer before adding the micelles.

### Vesicle Endocytosis

A 100 μL aliquot of the GUV suspension was carefully transferred into a 1.7 mL microcentrifuge tube (Fisher Scientific, USA). These vesicles were subsequently diluted 1:9 into a buffer consisting of 50 mM Na-bicine at pH 8.45, and 200 mM glucose to a final oleic acid concentration of 0.5 mM, enabling good contrast of vesicles against the background when imaged under fluorescence microscopy. This diluted vesicle solution was then split into ten 100 μL aliquots. Vesicle endocytosis was initiated by adding micelles from a 100 mM stock solution and Na-bicine buffer from a 1 M stock solution to a 100 μL diluted aliquot, then mixing by inverting the tube for ∼5 s. The added volumes and concentrations corresponding to each condition are summarized and listed in Table 1. Vesicle suspensions were allowed to equilibrate for at least 1 h before microscopy.

### Washing Endocytosed Vesicles

To dilute the free (unencapsulated) aqueous dye for imaging after passive endocytosis, the vesicles were diluted 1:9 into a 200 mM glucose buffer that was identical in composition and pH to the original buffer but lacked the lipids and dye. After centrifugation at 2000 g for 30 s, the top 90% of volume was removed by pipetting. The remaining solution was then agitated to resuspend the vesicles.

### Sequential endocytosis experiments

150 μL of the washed vesicle suspension was transferred into one well of a Nunc Lab-Tek II 8-well chambered coverslip (Thermo Scientific, USA) and the sample allowed to settle for 10 minutes to form a GUV monolayer at the bottom of the chamber. The endocytosis triggers were then pipetted into the open well in the sequence described in Figure 3.

Specifically, the initial GUVs that encapsulated calcein blue (blue) were prepared following the protocol outlined in the “Preparation of Micelles and Giant Unilamellar Vesicles (GUVs)” section. After diluting the GUVs 1:9 into a buffer containing 50 mM Na-bicine at pH 8.45 and 200 mM glucose, to a final oleic acid concentration of 0.5 mM, the GUVs were allowed to settle for 10 minutes. Subsequently, the first endocytosis trigger was added to the well, resulting in a final concentration 1 μM of 5’-Cy5-C_(10)_ A_(18)_-3’ along with ΔC_V_ = 100 mM Na-bicine and ΔC_A_ = 2.5 mM oleate. Inner compartments containing encapsulated 5’-Cy5-C_(10)_ A_(18)_-3’ (magenta) derived from the external solution were formed (see also Movie S1). After one hour, a second round of passive endocytosis was initiated by pipetting in a second stimulus to a final concentration of 5 μM of Fluorescein-12-UTP, ΔC_V_ = 100 mM Na-bicine and ΔC_A_ = 2.5 mM oleate. The changes in Na-bicine and oleate concentration were relative to the end of the first endocytic event; the final Na-bicine concentration was 250 mM, and the final oleate concentration was 5.5 mM. This second trigger led to the formation of new inner compartments containing encapsulated Fluorescein-12-UTP (yellow) derived from the external solution.

### Confocal Microscopy

Confocal images were collected using a Nikon A1R HD25 confocal laser scanning microscope equipped with LU-N4/N4S 4-laser unit.

### Vesicle Endocytosis Videos S3-S8

An aliquot of the GUV suspension encapsulating 1 mM HPTS was diluted 1:9 into a buffer consisting of 50 mM Na-bicine, pH 8.45, and 200 mM glucose to a final oleic acid concentration of 0.5 mM. 150 μL of the diluted suspension was transferred into one well of a Nunc Lab-Tek II 8-well chambered coverslip (Thermo Scientific, USA) and the sample allowed to settle for 10 minutes to form a GUV monolayer at the bottom of the chamber. 2 μL of a solution containing both 500 mM Na-bicine and 25 mM oleate micelles was then pipetted into the well containing the diluted GUVs. Epifluorescence microscopy was performed using a Nikon TE2000-U inverted microscope. A blue LED was used to excite the sample using a CoolLED pE-300^ultra^ system, and images captured using a pco.edge 4.2 sCMOS camera.

### Immersed boundary method simulations

To simulate growing, permeable vesicles under osmotic shock, we used the immersed boundary method to simulate the coupled fluid-structure interaction of permeable vesicles growing in an incompressible aqueous solution.^26^ Given the complexity of possible changes in vesicle morphology, it was important to perform simulations in three dimensions, and for computational efficiency we used the method described in Fai *et al*..^27^

For simplicity, we held the inner and outer osmolyte concentrations constant and did not account for solute exchange across the membrane (i.e. we set *P*_*s*_ = 0 in the notation of Sacerdote *et al*.).^24^ Given that the rate of solute exchange is typically much lower than that of solvent, this approximation is expected to be reasonable for the relatively short timescales of interest here. Further details can be found in SI Appendix I.

