## Supplemental Information for "Passive endocytosis in model protocells"

### **Supplementary Information for**
Passive endocytosis in model protocells

Stephanie J. Zhang[1,2], Lauren A. Lowe[3,4,5], Palapuravan Anees[6], Yamuna Krishnan[6,7,8], Thomas G. Fai[9]*, Jack W. Szostak[1,2,8,10]*, and Anna Wang[3,4,5]*

[1] Department of Chemistry and Chemical Biology, Harvard University, 12 Oxford Street, Cambridge, Massachusetts 02138, United States;
[2] Department of Molecular Biology, Center for Computational and Integrative Biology, Massachusetts General Hospital, Boston, MA 02114, United States;
[3] School of Chemistry, UNSW Sydney, Bedegal Country, New South Wales 2052, Australia
[4] Australian Centre for Astrobiology, UNSW Sydney, Bedegal Country, New South Wales 2052, Australia
[5] ARC Centre of Excellence in Synthetic Biology, UNSW Sydney, Bedegal Country, New South Wales 2052, Australia
[6] Neuroscience Institute, University of Chicago, Chicago, IL 60637, United States;
[7] Institute of Biophysical Dynamics, University of Chicago, Chicago, IL 60637, United States;
[8] Department of Chemistry, University of Chicago, Chicago, IL 60637, United States;
[9] Department of Mathematics, Brandeis University, Waltham, Massachusetts 02453, United States;
[10] Howard Hughes Medical Institute, Massachusetts General Hospital, Boston, MA 02114, United States;

*co-corresponding authors: Thomas G. Fai, Jack W. Szostak, and Anna Wang

Present address: Stephanie J. Zhang, Department of Pathology, Brigham and Women's Hospital, Harvard Medical School, 60 Fenwood Rd, Boston, Massachusetts 02115, USA.

**This PDF file includes:**

Figures S1 to S6
Movies S1 to S12
Appendix I

**Calculations for surface area increase**
Here we seek to estimate an upper bound for change in surface area that occurs during a topological change, which usually completes within seconds (see Movie S1, Movies S3-S8). The presumed growth process is that described by Chen and coworkers: an initial fast micelle growth phase leads to a 40% increase in surface area within 50 seconds, with further growth occurring via a different mechanism over minutes.[28]

To estimate the increase in surface area from the addition of micelles, we analyze the vesicle sizes before and after micelle addition. Figure 2 shows that the addition of 10 equivalents of micelles and a 25 mM osmotic shock results in ~15% of vesicles having at least 3 compartments. For such a case, assuming the compartment radius $(r)$ is half that of the external membrane (radius $R$), the final volume of the lumen is $\frac{4}{3}\pi R^3 - 3\frac{4}{3}\pi r^3 = \frac{4}{3}\pi R^3 - 4\pi(\frac{R}{2})^3 = \frac{5}{6}\pi R^3$. The lumen volume should have decreased by a factor of 2/3 following the osmotic shock, thus the original volume of the vesicle prior to the osmotic shock and micelle addition is $\frac{15}{12}\pi R^3$. Because the starting vesicle was a sphere, the initial radius is given by $(\frac{\frac{15}{12}\pi R^3}{\frac{4}{3}\pi})^{1/3} \sim 0.979\ R$. The initial surface area is thus $4\pi R^2 \cdot 0.979^2$. Comparing this to a final surface area of $4\pi R^2 + 4\pi(\frac{R}{2})^2 \cdot 3 = 7\pi R^2$ reveals a surface area increase of 1.8-fold. This is in agreement with the results from Zhu *et al.*, which found a doubling in surface area over five minutes upon the addition of 5 equivalents of micelles to multilamellar vesicles.[8] The increase in surface area over the timescales of the endocytic event are therefore expected to be no more than two-fold.

**Figure S1.** Two-dimensional schematic of topological transformations from a GUV into a stomatocyte shape (a), or a vesicle-in-vesicle structure (b).

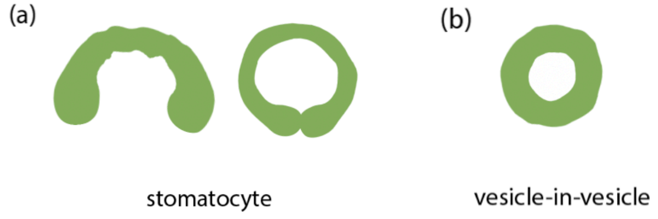

**Figure S2.** Whether an internal bud is connected to or disconnected from the outside can be distinguished by the addition of an impermeable aqueous dye. Schematics showing the outcomes of adding such a dye are shown in (a) and (b). (c) Confocal micrographs of vesicles containing calcein blue (blue) and endocytic compartments, after the addition of aqueous dye Alexa 594 hydrazide (red). The separate channels and merged image confirm that the internal compartments are disconnected from the outside. Scale bars represent 5 μm.

Experimental conditions: A GUV solution consisting of 5 mM of oleic acid encapsulating 1 mM calcein blue were prepared as described in the Materials and Methods section. The resulting solution was diluted 1:9 in a buffer composed of 50 mM Na-bicine (pH 8.45) and 200 mM glucose to obtain a final oleic acid concentration of 0.5 mM. Micelles from a 100 mM stock solution and Na-bicine buffer from a 1 M stock solution were added to 100 µL of the diluted vesicle solution, where $\Delta C_V$ = 100 mM Na-bicine and $\Delta C_A$ = 1 mM oleate. After allowing the vesicle suspensions to equilibrate for at least 1 hour, Alexa 594 hydrazide was introduced to a final concentration of 10 µM. The suspensions were mixed by inverting the tube for approximately 5 seconds before microscopy.

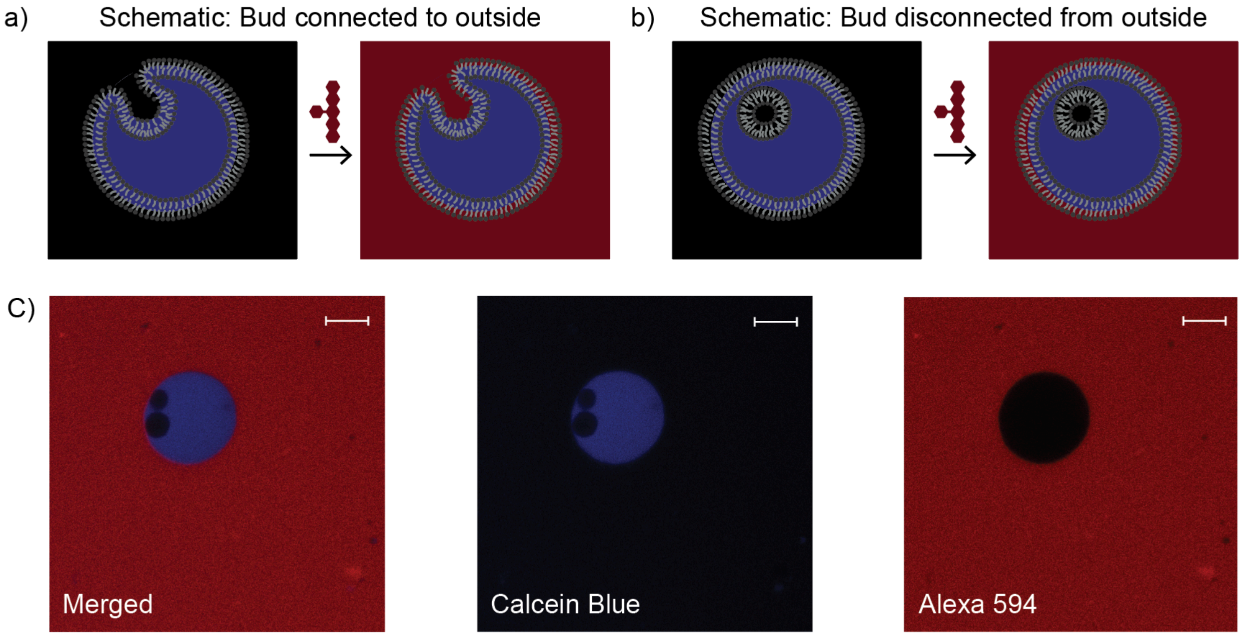

**Figure S3**. Whether an internal bud is connected to or disconnected from the external membrane can be distinguished by the addition of an impermeable membrane dye. Schematics showing the consequence of adding the membrane dye Voltair$^{PM}$ (yellow) to the vesicles are shown in (a) and (b). (c) Confocal micrographs of vesicles containing calcein blue (teal) and endocytic compartments, after the addition of the membrane dye Voltair$^{PM}$ (yellow). The separate channels and merged image confirm that the internal compartments are disconnected from the outside. Scale bars represent 5 μm.

Experimental conditions: A GUV solution consisting of 5 mM of oleic acid encapsulating 1 mM calcein blue were prepared as described in the Materials and Methods section and subsequently diluted 1:9 into a buffer containing 50 mM Na-bicine (pH 8.45) and 200 mM glucose, resulting in a final oleic acid concentration of 0.5 mM. Micelles from a 100 mM stock solution and Na-bicine buffer from a 1 M stock solution were added to the diluted vesicle solution, causing a change in concentration denoted as $\Delta C_V$ = 100 mM Na-bicine and $\Delta C_A$ = 1 mM oleate. Following at least 1 hour of equilibration, Voltair$^{PM}$ was introduced to a final concentration of 250 nM. The suspensions were mixed by inverting the tube for approximately 5 seconds before microscopy.

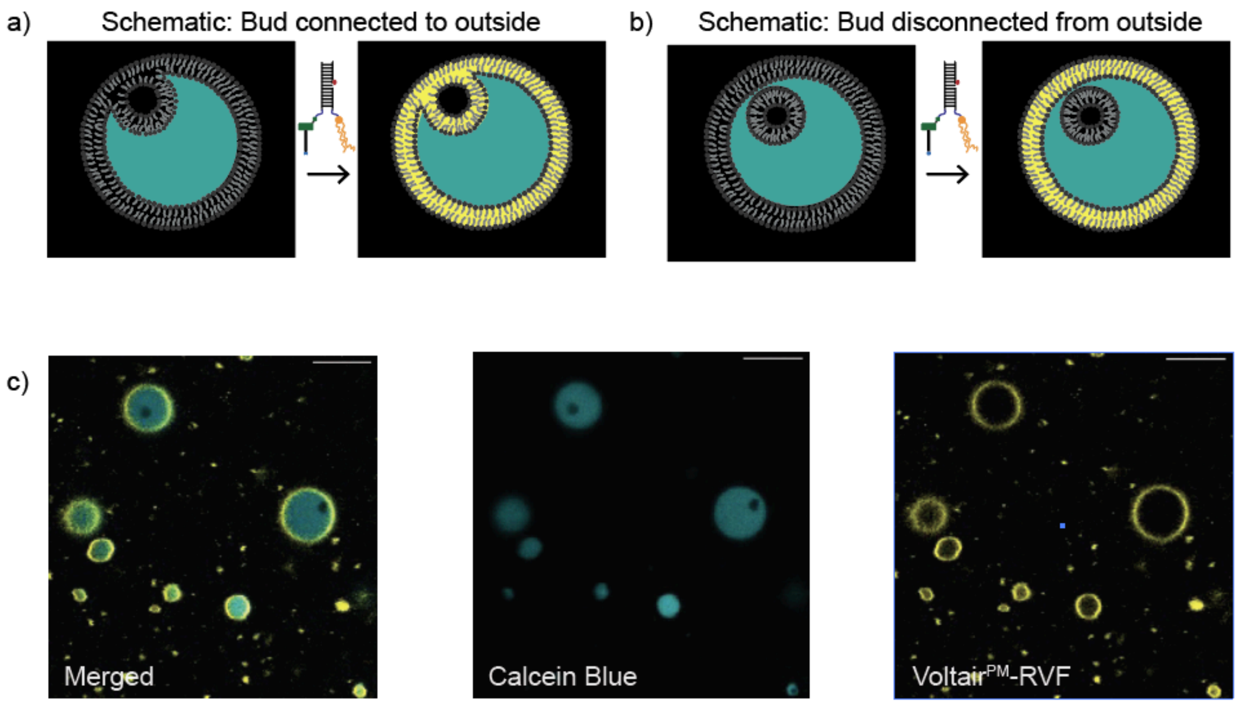

**Figure S4**. To calculate the average time $T_1$ required for a single molecule to enter a 4-$\mu\text{m}$-diameter vesicle, Equation (3) is rearranged and $dN$ is set to 1, giving $T_1 = 1/\,(P \cdot \Delta C \cdot A)$. This is calculated for a range of permeabilities spanning values for common nutrients ($P$ ranging from $10^{-10}$ to $10^{-13}$ cm/s), and a $\Delta C$ = 1 mM nutrient pool.

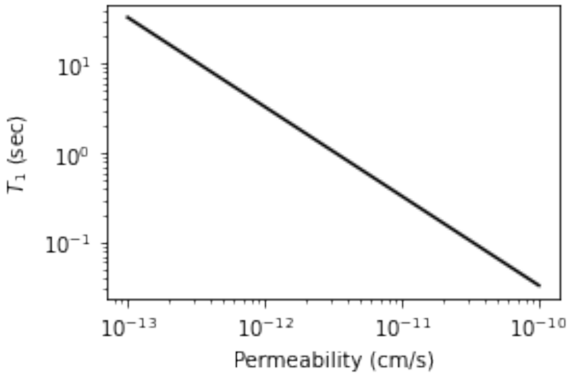

**Figure S5.** Internal compartments can slowly deliver nutrients of low permeability to the main lumen or 'cytoplasm' of a protocell. (a) 1 mM of HPTS, a relatively impermeable substrate, is encapsulated in the interior compartment via one hyperosmotic shock coupled with micelle addition ($\Delta C_V$ = 100 mM Na-bicine, $\Delta C_A$ = 2.5 mM oleate). (b) The ratio of fluorescence intensity of the internal compartment vs the 'cytoplasm' decreases with time; similarly, the ratio of fluorescence intensity of the internal compartment vs the external medium decreases with time, indicating that the HPTS inside the internal compartment is slowly entering the 'cytoplasm'.

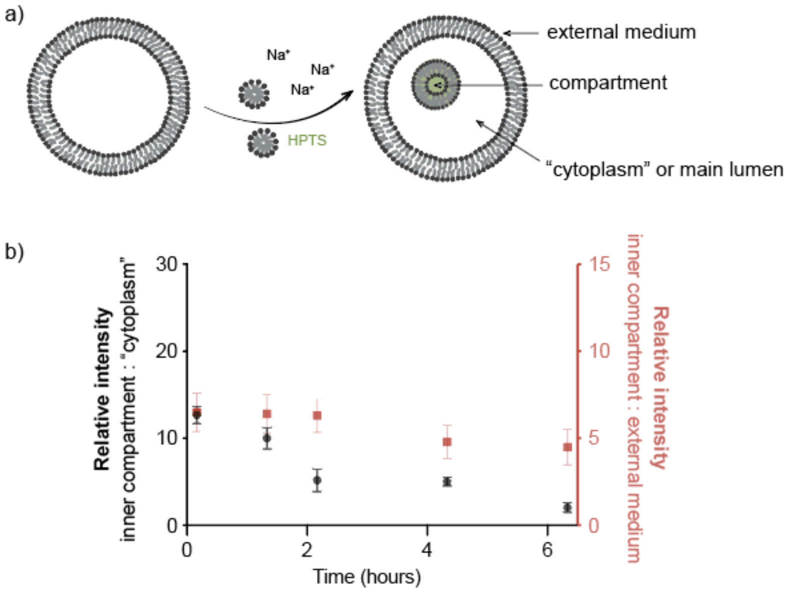


**Figure S6**. The release and retention of molecules with permeabilities of $10^{-12}$ and $10^{-13}$ cm/s across the membranes as a function of time is calculated by integrating Equation (3). (a) The intersection of the black dashed line (*y* = 0.9) and the black solid line indicates that 90% of the molecules with *P* = $10^{-12}$ cm/s will enter the main lumen of the protocell from the compartment within 11 hours. (b) The intersection of the black dashed line (*y* = 0.1) and the black solid line indicates that 10% of the molecules with *P* = $10^{-13}$ cm/s will enter the main lumen of the protocell from the compartment within 5 hours.

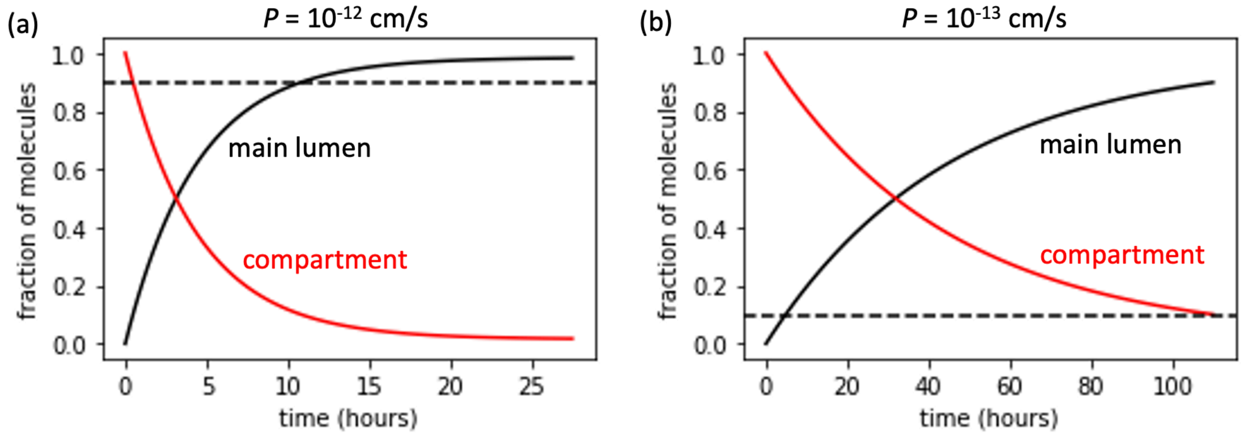

**Movie S1**. This real-time movie shows the topological transformation (prolate–oblate–stomatocyte-sphere) upon an osmotic shock and micelle addition (stimulus A; $\Delta C_V$ = 100 mM, $\Delta C_A$ = 1 mM). Prolate spheroid vesicles appeared to keep their shape for ~1 s, and then transformed into an oblate spheroid then stomatocyte within the next ~2 s. Subsequently, stomatocytes transformed into sphere-in-sphere structures via further invagination and pinching off of the internal vesicles. Each micrograph is 126 μm by 126 μm.

**Movie S2**. 3D reconstruction of vesicle-in-vesicle structures from sectional image slices along the *z*-axis, from slices taken using confocal microscopy.

**Movies S3-S8**. These real-time movies show the transformation (prolate–oblate–stomatocyte-sphere) upon an osmotic shock and micelle addition delivered by pipetting a small aliquot of concentrated micelles and buffer in the vicinity of the vesicles (see Methods). Depending on the proximity of the pipetted stimulus, the vesicles formed either one or more internal compartments. Each micrograph is 111 μm by 111 μm unless otherwise indicated.

**Movies S9-S11**. These movies show the simulated transformation (prolate–oblate–stomatocyte) upon an osmotic shock and micelle addition. Depending on the values of parameters used (see Table 2), in particular the bending modulus $K_b$ and the spontaneous curvature $c_0$, the vesicles formed either one or more internal compartments.

**Movie S12**. This movie shows that population diversity can be achieved from a uniform starting solution of vesicles, because of multiple heterogeneous stimuli (conditions as per Figure 3).

#### APPENDIX I

##### IMMERSED BOUANDARY SIMULATIONS

###### A. Formulation

In order to simulate the dynamics of growing fatty acid vesicles immersed in fluid, we use the immersed boundary method previously described in [1]. This method allows us to compute the emergent deformations of the vesicle as a function of the growth rate. We highlight the key aspects of this method here for the readers' convenience, following closely the description in [1].

The vesicle is modeled as an elastic shell having bending rigidity $B$, spontaneous curvature $c_0$, and local stretching resistance $k_a$, and the fluid surrounding the vesicle is modeled explicitly by the incompressible Navier-Stokes equations with fluid velocity $\mathbf{u}$ and pressure $p$. The fluid-structure interaction is specified with the vesicle parameterized in Lagrangian coordinates $\mathbf{q} = (q,r)$ and the vesicle configuration $\mathbf{X}(\mathbf{q},t)$ in cartesian coordinates. At any instant in time $t$, the vesicle configuration $\mathbf{X}(\mathbf{q},t)$ determines the elastic force density $\mathbf{F}(\mathbf{q},t)$ exerted on the fluid. In addition, the elastic material moves at a velocity

$$\frac{\partial \mathbf{X}}{\partial t}(\mathbf{q},t) = \mathbf{U}(\mathbf{q},t) + K\left(\Delta\phi + \frac{(\mathbf{F}(\mathbf{q},t)\cdot\mathbf{N}(\mathbf{q},t))}{\left\|\frac{\partial \mathbf{X}}{\partial r}\times\frac{\partial \mathbf{X}}{\partial s}\right\|}\right)\mathbf{N}(\mathbf{q},t), \tag{S.1}$$

where $\mathbf{U}(\mathbf{q},t) = \mathbf{u}(\mathbf{X}(\mathbf{q},t),t)$ is the fluid velocity at the vesicle membrane, $\mathbf{N}$ is the unit surface normal, $\Delta\varphi = (c_{\text{in}} - c_{\text{out}})k_B T$ is the osmotic pressure drop, and $K$ is the vesicle permeability. The velocity above is similar to those used previously for porous membranes [2–4], here with an additional osmotic pressure term. It is equivalent to specifying the local flux across the membrane to be proportional to the pressure jump [5].

Vesicle growth is realized through increasing the vesicle surface area over time uniformly over the surface. In particular, the vesicle is assumed to incorporate additional lipid amphiphiles at a constant rate $\gamma$ per unit area, so that $\mathrm{d}A/\mathrm{d}t = \gamma A$. On the other hand, changes in volume are determined implicitly through (S.1). In the limit of an impermeable membrane with $K = 0$, the vesicle moves at the local velocity of the incompressible fluid so that the enclosed volume is conserved. For nonzero $K > 0$, however, the membrane porosity causes the enclosed volume to change over time. In particular, in the absence of membrane forces ($\mathbf{F} \equiv 0$), we have:

$$\mathrm{d}V/\mathrm{d}t = KA\Delta\varphi, \tag{S.2}$$

so that osmotic shocks lead to vesicle swelling or shrinking. However, since the vesicle is assumed to be elastic, we have $\mathbf{F} \neq 0$ in general, and the changes in volume are the result of both osmotic pressures and elastic stresses on the membrane.

Remeshing is done periodically to keep the triangles in the mesh as regular as possible through growth and deformation, following the remeshing operations described in [6]. To avoid dealing with the changing topologies (which is nontrivial from both the modeling and numerical perspectives), simulations are halted prior to any budding or vesiculation events.

---

[6] M. Botsch, L. Kobbelt, M. Pauly, P. Alliez, B. Levy, L. Kobbelt, M. Pauly, P. Alliez, and B. Levy, Polygon Mesh Processing. A K Peters/CRC Press, 2010.
